# An ER–Inner Membrane Complex Bridge via TgVAP-TgVPS13A-TgDAT1 Drives Daughter Budding in *Toxoplasma gondii*

**DOI:** 10.64898/2026.01.02.697355

**Authors:** Lin Zhao, Jiawen Fu, Swaroop Peddiraju, Heming Chen, Keqin Huang, Ying Zhang, Nishith Gupta, Qian Jiang, Honglin Jia

## Abstract

All alveolates including apicomplexa parasites contain an inner membrane complex (IMC) underneath the plasma membrane. The IMC is synthesized *de novo* during the daughter cell budding (endodyogeny) within the mother cell and serves as a crucial scaffold for supporting cytoskeletal structures and the glideosome machinery for the parasite locomotion. However, the mechanism(s) underlying the membrane biogenesis in the IMC are not well understood. Using clinically-relevant and globally-prevalent pathogenic protist model, *Toxoplasma gondii*, we identified the TgVAP-TgVPS13A-TgDAT1 complex bridging the IMC to the endoplasmic reticulum (ER) – the major site of phospholipid synthesis. Individual components of this complex play a crucial role in the IMC biogenesis, where the multi-modular TgVPS13A protein interacts with the lipid scramblase TgDAT1 in the IMC *via* its C-terminal VAB domain, and with the ER-resident TgVAP through its N-terminal region. DAT1 is recruited for the progeny formation sites during the early stages of budding. Conditional depletion of TgVPS13A, TgDAT1 or TgVAP results in collapse of the inner membrane complex, leading to parasite death, as visualized by endodyogeny-specific organelle markers. LactC2-GFP, a biosensor of phosphatidylserine and phosphatidylthreonine lipids made in the ER and enriched in the IMC, also mislocalizes upon protein depletion. In conclusion, we propose that TgVAP-TgVPS13A-TgDAT1 bridge the ER and IMC and mediate the inter-organelle transport of lipids, thus contributing to the organelle biogenesis and daughter budding in *T. gondii*.

**Author Summary:** The inner membrane complex (IMC) in apicomplexan parasites is essential for maintaining the structural stability of the parasite, as well as for its budding and motility. The IMC is composed of flattened vesicles, alveolins, and microtubules. However, the mechanism behind the biogenesis of these flattened vesicles remains unclear. In this study, we provide evidence that a protein complex formed by TgVAP, TgVPS13A, and TgDAT1 exists between the endoplasmic reticulum (ER) and the IMC. Phenotypic analysis indicates that the absence of any individual component of this protein complex disrupts the biogenesis of daughter IMCs, which in turn affects the budding of daughter parasites. We also demonstrated that TgDAT1 functions as a scramblase. Based on our findings, we propose a model in which the TgVAP-TgVPS13A-TgDAT1 complex mediates lipid transport from the ER to the nascent IMC, driving the expansion of the IMC membrane and the budding of daughter parasites in *T. gondii*. Furthermore, TgDAT1 may facilitate the transfer of lipids from the outer leaflet to the inner leaflet of the IMC sacs, promoting a balanced composition of the IMC membrane components.

## Introduction

Apicomplexa are obligate intracellular parasites that pose significant health risks to humans and animals. Alongside ciliates and dinoflagellates, these protists belong to the superphylum Alveolata, which is hallmarked by a unique structure comprising membrane vesicles (alveoli, cisternae) situated just underneath the plasma membrane. In Apicomplexa, this membrane structure is referred to as the inner membrane complex (IMC). The IMC together with the plasma membrane forms the pellicle of the parasites, which harbors the actin-myosin motors and other locomotion-related protein machinery cumulatively termed as Glideosome. In *T. gondii*, the IMC is made up of flattened vesicles, networks of alveolin, and sub-membranous microtubules. This cytoskeletal structure is essential for the parasite’s replication, movement, and invasion of host cells(1). The IMC begins from an apical polar ring (APR), and the apical region above the APR is encapsulated by the plasma membrane. The upper part of the IMC consists of a single conical IMC vesicle, while the structure at the base is known as the basal cap. Vesicles within the IMC are connected by longitudinal and transverse suture proteins (2). During the progeny budding (endodyogeny), the IMC is made *de novo*. However, little is known about the mechanisms that govern the biosynthesis of the flattened vesicles within the IMC.

Membrane expansion needs a continuous supply of bulk lipids and transport from the source organelles, such as ER or lipid droplets (LDs). In other eukaryotes, lipid transfer proteins (LTPs) are proven to accomplish this process at the membrane contact sites (MCSs), where two organelles are proximal so that a single protein can span the entire distance between membranes to transfer lipids (3). MCSs are considered to play a central role in non-vesicular lipid transport. Currently, ten members of LTPs have been identified, including VPS13 (A-D), ATG2 (A-B) and BLTPs (1, 2, 3A-B) (4). VPS13 is a new class of LTPs that emerged recently. While several species, including plants and protists, contain multiple VPS13 genes, yeast possesses a single VPS13 protein. This family of proteins appears at multiple MCSs to mediate lipid transport in a way similar to ATG2 by forming a hydrophobic channel at MCSs (5–8). Fragments of VPS13 were shown to bind multiple lipids and to transfer lipids between artificial liposomes (7). For instance, human VPS13A resides at the MCSs between ER and mitochondria, endosome/lysosomes, lipid droplet (LDs) (7, 9), or plasma membrane (10), VPS13B is localized at the interface of Golgi cisternae (11) or between ER exit site (ERES) and the Golgi (12), and VPS13C is localized at the MCS of the ER and endo/lysosomes (13), whereas VPS13D localizes at between the ER and mitochondria or peroxisomes (14).

The N-terminal region of VPS13 proteins is responsible for binding to adaptors at the donor organelle, which supply lipids. In contrast, the adaptors located at the opposing membrane interact with either the VAB domain or the C-terminal region of VPS13. For example, human VPS13 interacts with VAP at the endoplasmic reticulum (ER) through its N-terminal region (9, 13), whereas binds the scramblase XK at the plasma membrane through the VAB domain (10), VPS13B interacts with Sec23 at the ERES (12), VPS13C binds Rab7 at the endo/lysosomes (7), and VPS13D binds the ESCRT-III component TSG101 (15). Once lipids are delivered to the outer leaflet of the nascent membrane, certain scramblases transfer the lipids from the outer leaflet into the inner leaflet to facilitate the expansion of the membrane bilayers. Till now, several classes of scramblases have been identified at various membranes, including TMEM16, XK, TMEM41B, VMP1, ATG9, Class A GPCRs, and CLPTM1L. Among them, TMEM16, Class A GPCRs, and XK act as the scramblases in the plasma membrane to transfer lipids (10, 16, 17). TMEM41B, VMP1, and CLPTM1L function in the ER (18), and ATG9 is recruited to the isolated membrane to fulfill the role of scramblase in the autophagy pathway (19).

We and other research teams have demonstrated that Golgi-mediated vesicular transport is crucial for the biogenesis of the IMC in *T. gondii* (20–24). However, phospholipids are predominantly synthesized in the ER, Mitochondrion and Golgi Network of the parasite (25–32). Hence, it is reasonable to speculate that a pathway exists in these parasites that directly mediates the transport of bulk lipids from the sites of lipid synthesis to the IMC and elsewhere. Furthermore, an efficient continuous supply of bulk lipids may be required for the membrane expansion of the IMC during daughter cell budding. A previous study has also indicated that MCSs exist at the IMC (33), but whether they contribute to the lipid transport to the IMC and mechanisms underlying the formation of flattened vesicles remain completely unknown. This study identified an ortholog of VPS13 in *T. gondii* (TgVPS13A), which interacts with a VAMP-associated protein (VAP) in the ER (TgVAP) and with a major facilitator superfamily (MFS) transporter located in the IMC (TgDAT1). TgVPS13A, TgVAP and TgDAT1 are required for the biogenesis of the IMC and progeny formation. The entire protein complex forms an inter-organellar bridge to mediate lipid transport at the ER-IMC interface.

## Results

### Identification of VPS13 and VAP family proteins in *T. gondii*

VPS13 family proteins bridge lipid transport through binding with adaptors at the donor and the target organelles (Fig 1A). Our BLAST analysis of the parasite genome (www.ToxoDB.org) resulted in four *VPS13* genes in *T. gondii* (Fig 1B and Fig S1A). Further bioinformatic analysis indicated that VPS13 orthologs harbor domains of this family, including a chorein-N domain followed by an extended chorein region, VPS13 adaptor binding region (VAB), and α-helical region (called the ATG2-C domain due to its homology with a corresponding region in ATG2) and Pleckstrin Homology (PH) domain at the C terminus (Fig S1A). Phylogenetic analysis indicated a conservation of VPS13 family in other alveolates including *Chromera velia*, the nearest free-living photosynthetic ancestor of apicomplexan parasites (Fig 1C). By fusing a HA tag, we observed that TGGT1_291180 (TgVPS13A) is more closely associated with IMC1 during daughter budding compared to TGGT1_232080 and TGGT1_306020 (Fig 1D). Our structure modeling of TgVPS13A revealed a bridge-like central domain flanked by the N-terminal chorein and C-terminal VAB and ATG2-C domains, positioned for interaction with partner proteins in the donor and acceptor membranes (Fig S1B). The endoplasmic reticulum (ER) is the primary site where most phospholipid synthesis occurs. Next, we determined to detect the association of the TgVPS13A to the ER, we inserted an SmFP-HA tag at the C-terminal through the CRISPR/Cas9 method (Fig S1C). The integration of the tags was confirmed by diagnostic PCR (Fig S1D). The results indicated that partial signal of TgVPS13A-SmFP-HA merged with that of the ER marker EGFP-HDEL (Fig 1E).

**Fig 1.**
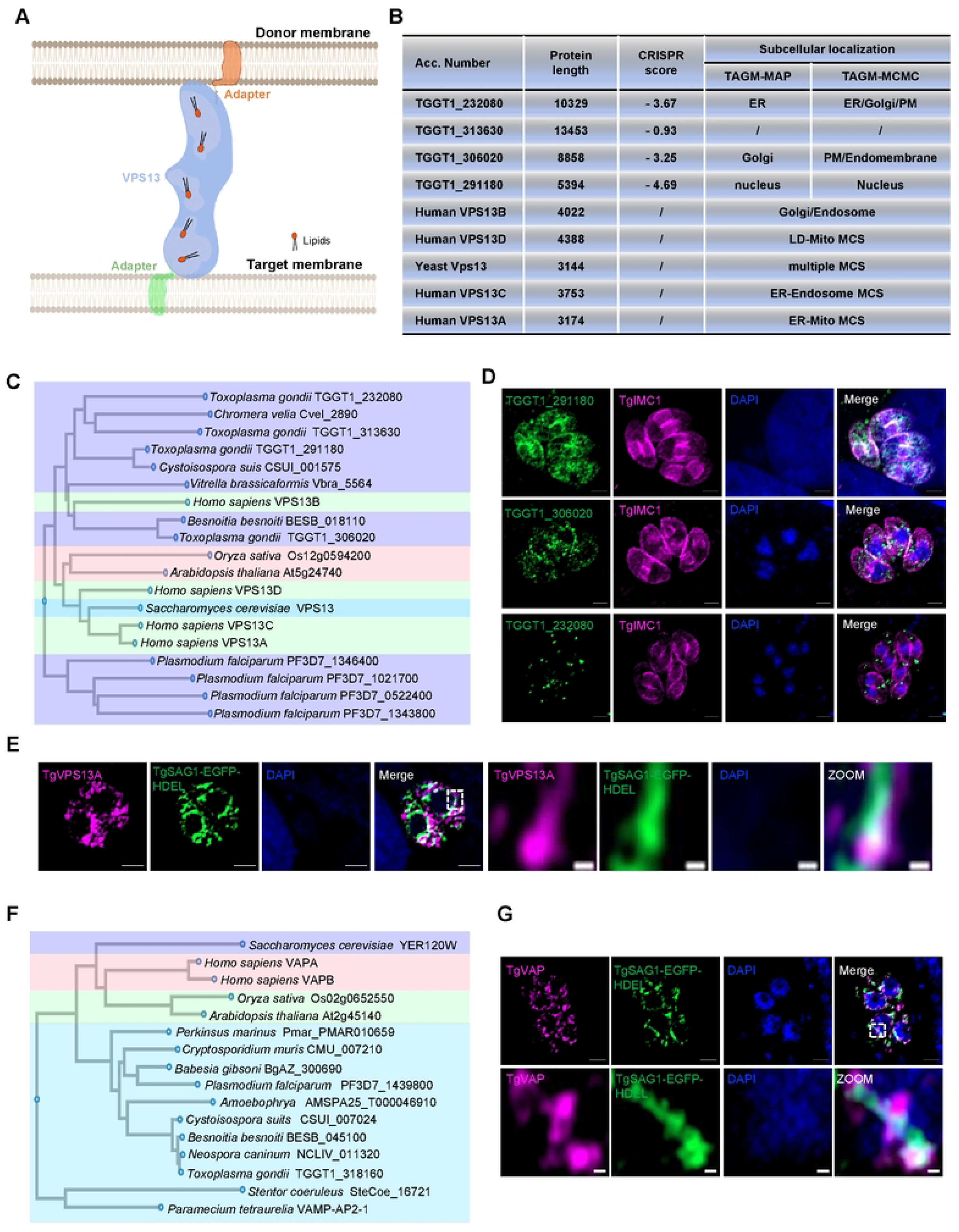
Identification of VPS13 and VAP family proteins in *T. gondii.* **(A)** A schematic diagram showing VPS13 family protein connecting donor and target organelles. **(B)** The list displays the length, score, and localization on donor and target organelles of VPS13 in *human*, *yeast*, and *T. gondii*. **(C)** Phylogenetic analysis of VPS13 by CLUSTALW. **(D)** IFA shows the localization of VPS13 proteins in *T. gondii*. **(E)** IFA shows that VPS13 is localized to the ER. **(F)** Phylogenetic analysis of TgVAP by CLUSTALW. **(G)** IFA showing colocalization of TgIMC1 and ER-resident proteins TgSAG1-EGFP-HDEL with TgVAP. Magenta: rabbit anti-HA antibodies; anti-TgIMC1 polyclonal antibodies; Green:Mouse anti-HA antibodies and EGFP signals; Blue: DAPI. Scale bars indicate 2 µm in the merged panels and 0.2 µm in the zoomed panels.

In mammalian cells, VPS13 anchors to the ER by binding integral membrane proteins known as VAMP-associated proteins (VAPs) through its N-terminal region, thereby facilitating lipid transport at MCSs (8, 12). Through a blast search using human VAPA and VAPB as baits, we found a unique VAP protein in *T. gondii* (TgVAP). TgVAP is phylogenetically conserved across alveolates, forming a distinct cluster separated from its human and yeast orthologs (Fig1 F). Sequence alignment of TgVAP with *Hs*VAP1/2 (Fig S2A) and *in silico* modeling displayed the occurrence of a conserved MSP domain for interaction with the FFAT motif of TgVPS13A (Fig S2B). Alphafold models of TgVPS13A N-terminus and TgVAP have been docked together using HADDOCK. The predicted complex reiterated the strong possibility of TgVAP binding to TgVPS13A N terminus as these models ranked higher in the cluster with the best scores (Fig S2C). We inserted an 6×HA tag at the N-terminal through the CRISPR/Cas9 method. Through an IFA analysis, we found 6×HA-TgVAP localized primarily in the ER (Fig 1G). The ER localization of TgVPS13A and TgVAP proteins is consistent with the hyperLOPIT database (37). The existence of the TgVAP and TgVPS13 complex at the ER-IMC interface was also evidenced in a proximity biotinylation assay, which was designed to screen novel IMC proteins, using the established IMC marker GAPM3 as a bait. In this analysis, both TgVPS13A and TgVAP were identified among the results (Fig S3A-E). Collectively, our data suggest the presence of a lipid transport complex formed by TgVPS13A and TgVAP between the parasite ER and IMC.

### TgVPS13A is partially associated with the nascent IMC

To ascertain the association of TgVPS13A with the IMC, we conducted its 3D colocalization analysis with TgIMC1. The fluorescence signal of the two proteins matches constantly over the distance at different stages of parasite development (Fig 2A-B). To monitor the dynamics of TgVPS13A signal during the emergence of daughter parasites, we colocalized TgVPS13A with TgIMC29 or TgGAPM3, marking the nascent progeny. TgVPS13A was indeed associated to the daughter cells during the budding (Fig 2C). In principle, both the ER of mother and daughter cells could contribute lipids needed for the budding IMC. We therefore examined TgVPS13A between the interface of the budding IMC and of the mother or daughter ER. To investigate this, we inserted an EGFP or a 3xV5 tag at the C-terminal of TgGAPM3 (budding IMC) and TgSEC61β (ER) (Fig 2D-E). TgSEC61β is an ortholog of the human SEC61β, which is commonly used as an ER marker (7). We confirmed its localization using an ER marker in *T. gondii* (Fig S3F). TgVPS13A was detectable at all inter-organelle interfaces.

**Fig 2.**
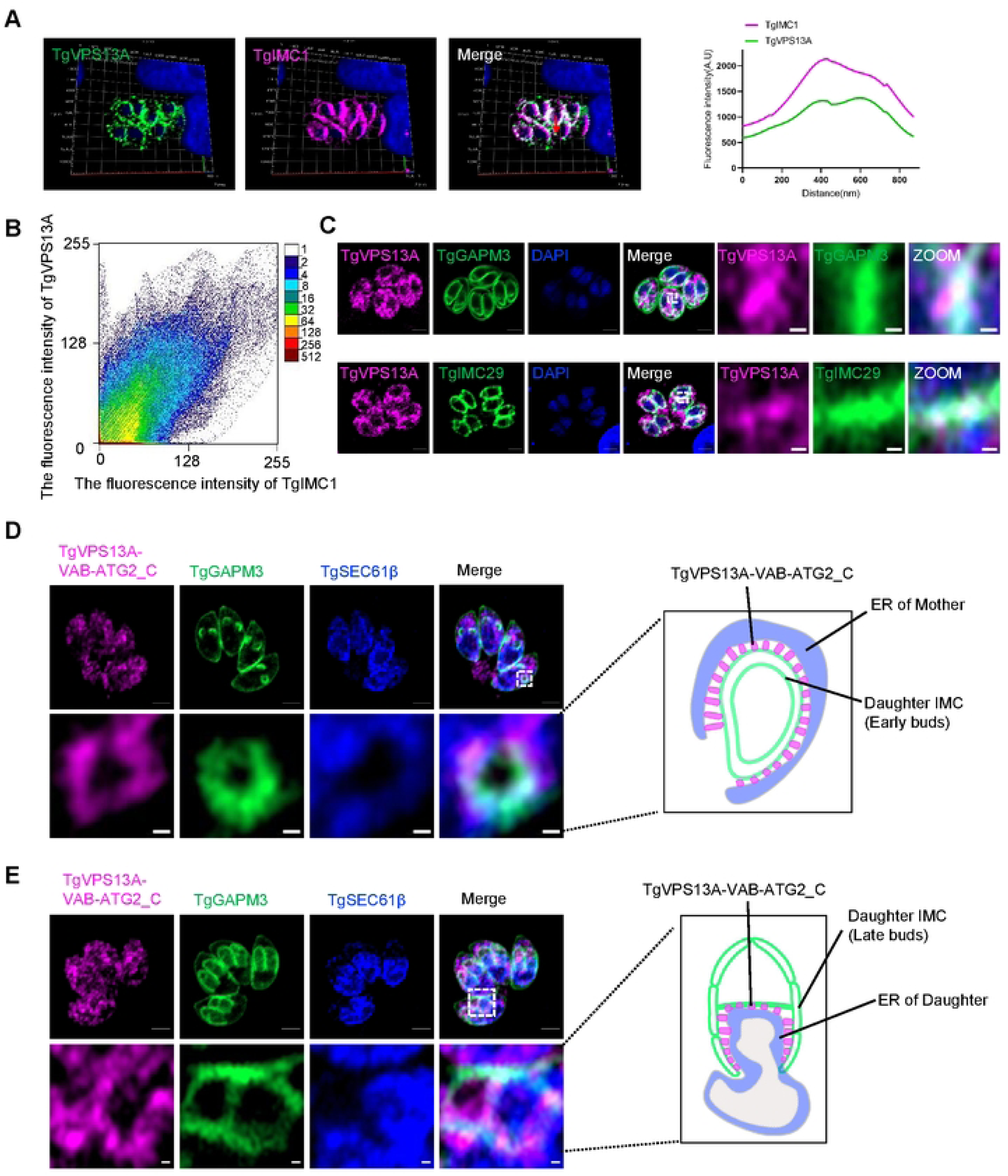
TgVPS13A is partially associated with the nascent IMC. **(A)** IFA showing a 3D colocalization of TgVPS13A-SmFP-HA with TgIMC1. The signal intensity of TgVPS13A-SmFP-HA across the TgIMC1 distribution are graphed. **(B)** Scatter plot of colocalization analysis of TgVPS13A and TgIMC1 using Image J. **(C)** Subcellular localization of TgVPS13A-SmFP-HA was investigated by co-staining with TgGAPM3 and TgIMC29. The EGFP tag was inserted at the C-terminus of TgIMC29 and TgGAPM3 in the parasites. **(D** and **E)** IFA showing the colocalization of TgVPS13A-VAB-ATG2-C with TgSEC61β and TgGAPM3. Magenta: anti-TgIMC1 polyclonal antibodies, rabbit anti-HA antibodies, mouse anti-Myc antibodies; Green: rabbit anti-HA antibodies and EGFP signals; Blue: DAPI, rabbit anti-V5 antibodies. Scale bars indicate 2 µm in the merged panels and 0.2 µm in the zoomed panels.

Our findings also demonstrated that the VAB domain, in conjunction with the ATG2-C region of TgVPS13A, is sufficient for its localization at the IMC (Fig 2D-E). Consequently, stable transgenic parasites expressing SmFP-MYC-tagged VAB-ATG2-C region of TgVPS13A were selected to examine the TgVPS13A expression at the interface between the ER and the IMC. The results indicated that the ER of the mother parasites interacts with the emerging buds of daughter IMCs during the initial phase. As the progeny develop, the ER of the budding parasites start associating with the middle region of the nascent IMC, but not the cap. It has been reported that the cap region develops distinctly compared to the bulk IMC (38), which is consistent with our datasets.

### TgVPS13A is essential for parasite survival and the IMC assembly

To investigate the physiological role of TgVPS13A in IMC biogenesis and parasite survival, we first attempted to regulate its expression by the AID and TATi systems (39, 40). However, these efforts were unsuccessful. Thus, we adapted the DiCre system for a conditional depletion of TgVPS13A (41). We inserted two loxP sites flanking the promoter region and the third exon of TgVPS13A in parasites expressing a rapamycin-inducible DiCre recombinase (Fig 3A). The correct insertion of the sequences was verified through diagnostic PCR and sequencing (Fig 3B). The expression of TgVPS13A remained unaffected in routine parasite cultures (-rapamycin). Upon treatment with rapamycin, the promoter region and the 5’-fragment of the *TgVPS13A* gene flanked by the loxP sites could be efficiently excised from the genomic DNA, resulting in the blockage of protein expression (Fig 3C). A depletion of TgVPS13A expression was confirmed by IFA (Fig 3C). The lytic cycle of the parasites was disrupted upon knockdown of TgVPS13A, as judged by the lack of plaque formation in host-cell monolayers treated with rapamycin (Fig 3D). Intracellular replication of the rapamycin-treated parasites was also compromised (Fig 3E).

**Fig 3.**
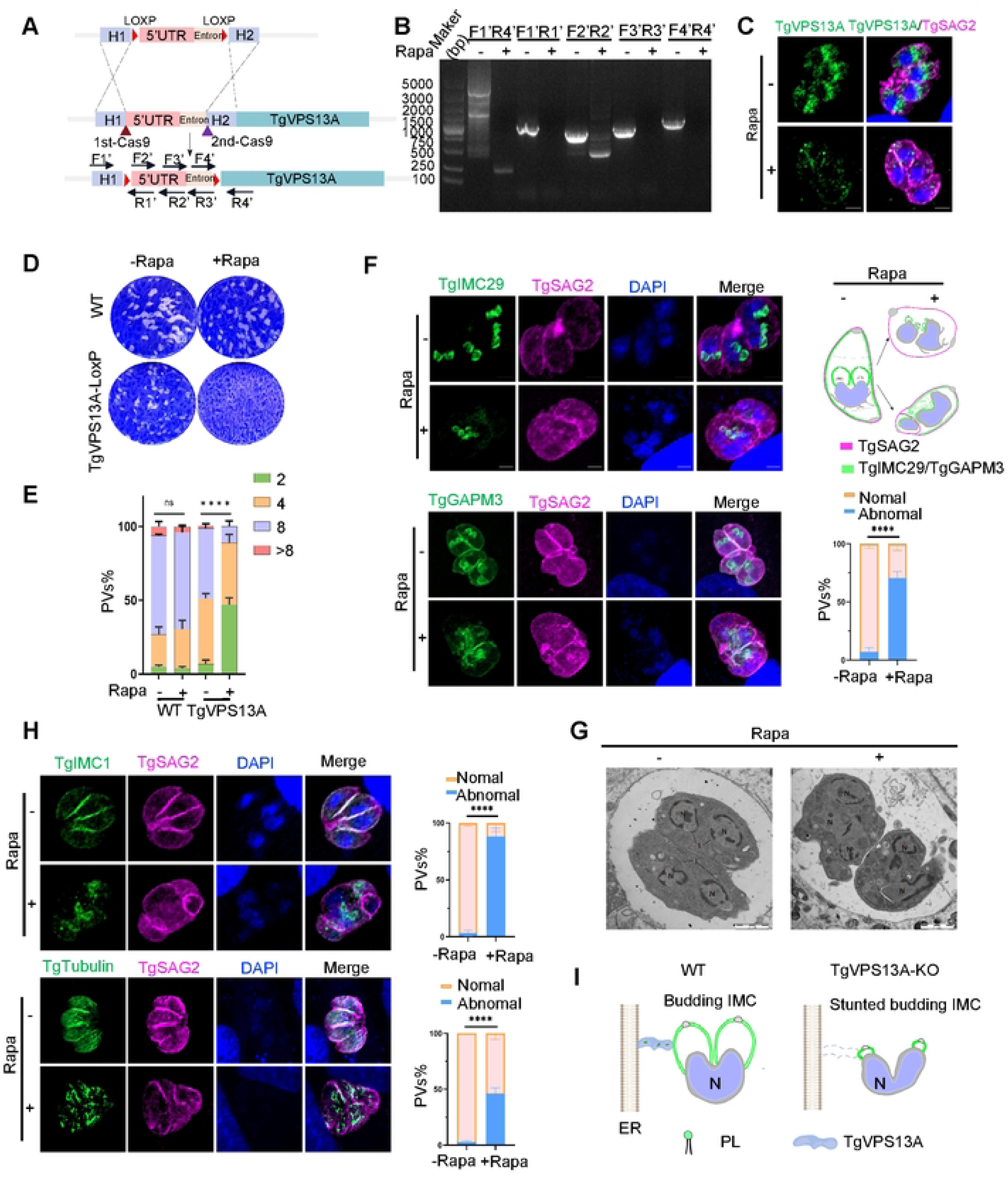
TgVPS13A is essential for parasite survival and is necessary for the assembly of the IMC. **(A)** The schematic representation illustrating the DiCre-LoxP system used to conditionally deplete TgVPS13A. **(B)** PCR analysis confirming the N-terminal recombination of TgVPS13A involving the two LoxP sites. **(C)** IFA analysis showing the expression of TgVPS13A in parasites cultured with or without rapamycin (Rapa) for 96 h. **(D)** Plaque assays were conducted by infecting BJ-5ta cells with the specified parasites for 9 days, either in the presence or absence of Rapa. **(E)** Replication of the indicated parasites in host cells was assessed after 96 h of culture with or without Rapa. The average number of parasites per vacuole was calculated, and the data were analyzed using a two-way ANOVA, revealing statistical significance (****P < 0.0001). **(F)** IFA analysis of IMC biogenesis involved staining for TgIMC29 and TgGAPM3. The abnormal and normal PVs were counted from TgGAPM3. The data were analyzed for statistical significance by the unpaired t-test; ****P<0.0001. Schematic diagram of the assembly of early buds of daughter IMCs in TgVPS13A-deficient parasites. **(G)** Transmission electron microscopy analysis of daughter parasites treated with and without Rapa for 96 h was performed. In the images, ‘N’ indicates the nucleus. **(H)** IFA analysis of the IMC-associated cytoskeleton by examining the localization of TgIMC1 and Tgtubulin in TgVPS13A-deficient parasites. The abnormal and normal PVs were counted from H. The data were analyzed for statistical significance by the unpaired t-test; ****P<0.0001. **(I)** A schematic model of the IMC biogenesis and daughter cell budding in TgVPS13A-delepleted parasites. PL indicates Phospholipid. Green: rabbit anti-HA antibody, rabbit anti-TgIMC1 polyclonal antibodies, rabbit anti-TgTubulin polyclonal antibodies, and EGFP signal; Magenta: rabbit and mouse anti-TgSAG2 polyclonal antibodies; Blue: DAPI. Scale bars represent 2 µm.

Our ensuing work investigated the assembly of the early buds by staining the daughter IMC proteins, TgIMC29 and TgGAPM3, in parasites lacking TgVPS13A. The absence of TgVPS13A led to collapsing of the nascent IMC and asynchronized budding, resulting in morphological abnormalities (Fig 3F). The cell division was disrupted in TgVPS13A-depleted parasites, as evident by the nuclear staining. We examined the cytokinesis process using transmission electron microscopy, which demonstrated a failure of cytokinesis (Fig 3G). Immunostaining of the IMC-associated cytoskeleton (TgIMC1 and TgTubulin) revealed similar defects in IMC budding and failure of tubulin assembly (Fig 3H). Taken together (schematized in Fig 3I), our data show that TgVPS13A is required for the IMC biogenesis and budding in *T. gondii*.

### TgVAP in the ER is required for the parasite survival and IMC biogenesis

As described, we found a VAP ortholog, located in the parasite ER. TgVAP also partially co-localized with TgVPS13A at the IMC both in mature and daughter parasites (Fig 4A). To test the interaction of TgVPS13A with TgVAP, we co-expressed the predicted N-terminal domain of TgVPS13A containing the FFAT motif and full length of TgVAP in HEK293T cells and performed co-immunoprecipitation. Our results showed that the 3xHA-tagged TgVPS13A-N_928-1392_ interacted with 3xFLAG-tagged TgVAP (Fig 4B). These results indicate that TgVAP acts as a adaptor for TgVPS13A on the ER, tightly anchoring TgVPS13A to the ER membrane. To investigate the role of TgVAP in the survival and assembly of the IMC in budding progeny, we inserted a 6xHA-AID* tag at the N-terminal of TgVAP, allowing its regulation by 3-indole acetic acid (IAA). The 5’-genomic integration of the tag was verified by diagnostic PCR (Fig 4C-D). Treatment with IAA induced efficient degradation of 6xHA-AID*-TgVAP in the parasites (Fig 4E-F). TgVAP-depleted parasites were compromised in their growth, as demonstrated by the lack of plaque formation in IAA-treated parasites cultured on host-cell monolayers (Fig 4G). The abnormal assembly of the IMC in the parasites was observed when TgVAP was depleted (Fig 4H), albeit these changes were not as severe as in TgVPS13A-deficient cells. Notably, the IMC-associated cytoskeleton (TgIMC1 and TgTubulin) was not affected in upon knockdown of TgVAP (Fig 4I). In brief, TgVAP anchored in the parasite ER is needed for the IMC biogenesis and budding (Fig 4J).

**Fig 4.**
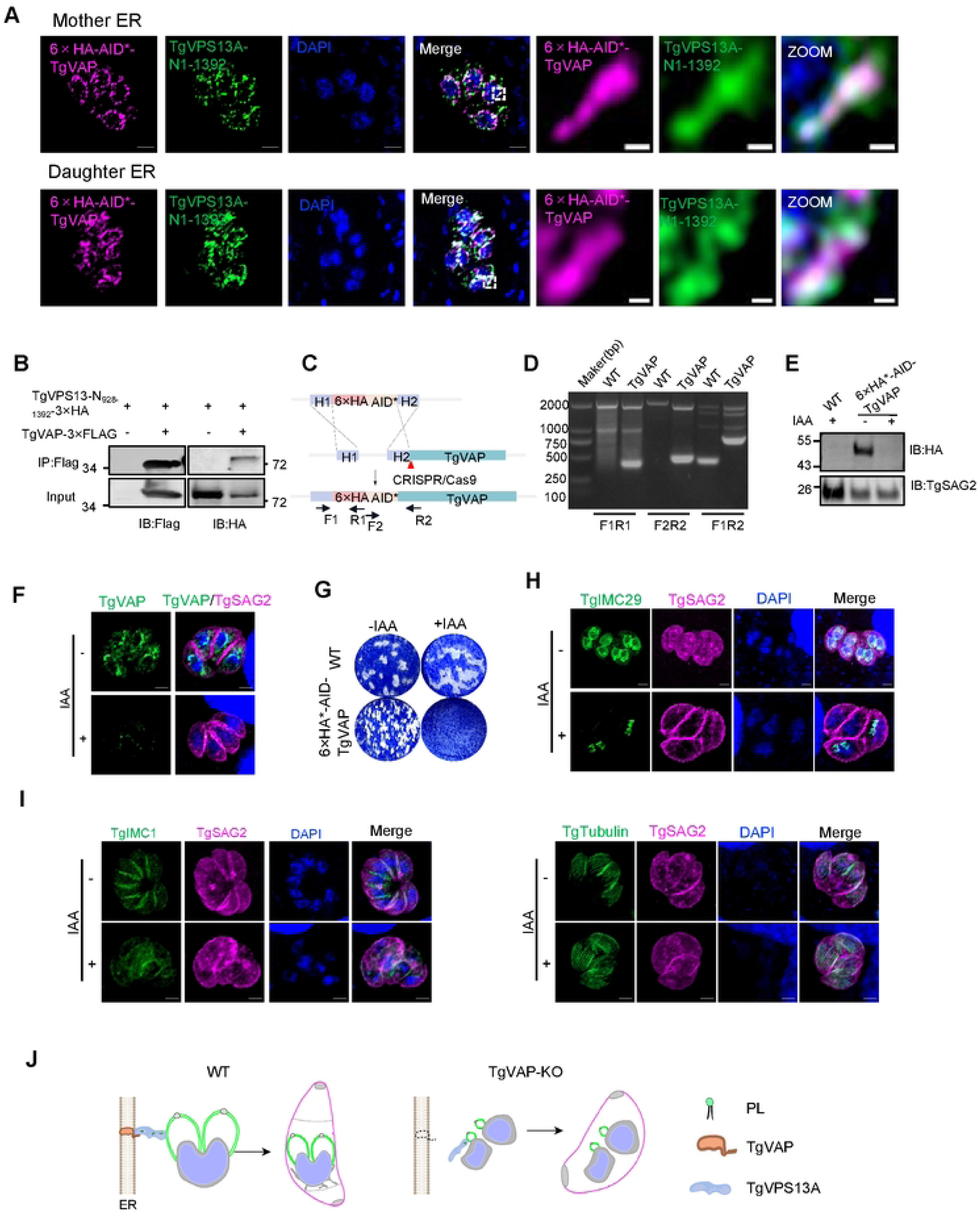
TgVPS13A is anchored to the ER by binding to TgVAP. **(A)** IFA was performed to examine the colocalization of TgVAP with TgVPS13A-N_1-1392_ before and after division. The parasites were transfected with a plasmid expressing TgTgVPS13A-N1-1392-SmFP-Myc, under the control of the native promoter. **(B)** Co-IP was performed using 3×FLAG-tagged TgVAP and 3×HA-tagged TgVPS13A-N928-1392aa. Cells were transiently transfected with plasmids expressing 3×FLAG-tagged TgVAP and 3×HA-tagged TgVPS13A-N928-1392aa. The resulting cell lysates were subjected to immunoprecipitation and then blotted with anti-FLAG and anti-HA antibodies. **(C)** A schematic diagram of inserting a 6×HA-AID* tag at the N-terminal of TgVAP. **(D)** PCR identification of the 6×HA-AID* tag inserted at the N-terminal of TgVAP. **(E)** The expression of TgVAP was examined using western blot analysis. The parasites were cultured in the presence of IAA for 24 h in BJ-5ta cells and then subjected to the analysis. WT refers to the wild type. TgVAP expression was detected using anti-HA antibodies. **(F)** IFA analysis of TgVAP expression in parasites with and without IAA treatment for 24 h. **(G)** Plaque assays were conducted by infecting BJ-5ta cells with the parasites for 9 days, both in the presence and absence of IAA. **(H)** The parasites were cultured with or without IAA for 48 h to assess the biogenesis of the IMC in daughter cells, using TgIMC29 for staining. The EGFP tag was inserted at the C-terminus of TgIMC29 in the parasites. **(I)** IFA analysis showing the impact of TgIMC1 and Tgtubulin in TgVAP-deficient parasites with and without IAA treatment for 48 h. **(J)** A schematic diagram of the effect of TgVAP on IMC biogenesis. Magenta: rabbit anti-HA antibody, mouse and rabbit anti-TgSAG2 polyclonal antibodies; Green: rabbit anti-HA antibody, mouse anti-Myc antibody, rabbit anti-TgTubulin polyclonal antibodies, EGFP signal and anti-TgIMC1 polyclonal antibodies. Scale bars measure 2 μm in the merged panels and 0.2 μm in the zoomed-in panels.

### TgDAT1 is a physiologically-essential IMC-localized lipid scramblase

A requirement of TgVPS13A-VAP proteins for the biogenesis of the nascent alveoli suggested the presence of a scramblase in the IMC vesicle membrane to bridge the transport of lipids (Fig 5A). Therefore, we performed a global screening by endogenously inserting a hemagglutinin (HA) tag using the CRISPR/Cas9 method (Fig S4A). We included the annotated transporters with more than two transmembrane domains from ToxoDB in this screening, excluding those with a CRISPR score greater than −1.0. In this screening, we detected fluorescence signals from 64 out of 71 annotated putative transporters encoded in the genome of *T. gondii* (Table S2 and Fig S4B). Our search identified TGGT1_258700 belonging to the major facilitator superfamily (MFS) of transporters, similar to other SPNS transporters (42, 43), resulting in localization of the fusion protein in the IMC (Fig 5B). The RNA expression profile of this transporter (termed as **D**aughter **A**lveoli **T**ransporter 1, DAT1) is similar to AC9, IMC29, and IMC32 (Fig 5C). An examination of the cellular localization of the transporter, through co-staining with the ISPs, revealed that this protein was predominantly expressed in early daughter buds but was rarely found in mature parasites (Fig 5D). Additionally, further co-localization analysis with the protein markers of different IMC regions revealed that its localization at the IMC during emerging process of daughter buds (Fig 5E-G). The Alphafold structure of TgDAT1 showed that its binding cavity is rich in hydrophobic residues, containing only a few charged residues (10/124) (Fig 5H). This suggests that its substrate likely has a lipidic/lipophilic nature. We then investigated whether TgDAT1 can transport lipids from the inner to the outer leaflet of liposomes using the previously established dithionite assay (19, 44). In this assay, liposomes containing fluorescent 7-nitro-2, 1, 3-benzoxadiazol (NBD) acyl-labeled phospholipids are treated with dithionite, a membrane-impermeable reducing agent. Dithionite reduces and irreversibly quenches the fluorescence of NBD-lipids located in the outer leaflet of the liposome (Fig 5I). TgDAT1 was expressed in HEK293T cells and purified (45) for this experiment. After detecting the fluorescence of NBD-PC for 50 seconds, dithionite was added, and measurements continued for an additional 600 seconds. The results showed that the fluorescence curve of TgDAT1 significantly dropped compared to the controls (Fig 5I), indicating that TgDAT1 acted as a scramblase.

**Fig 5.**
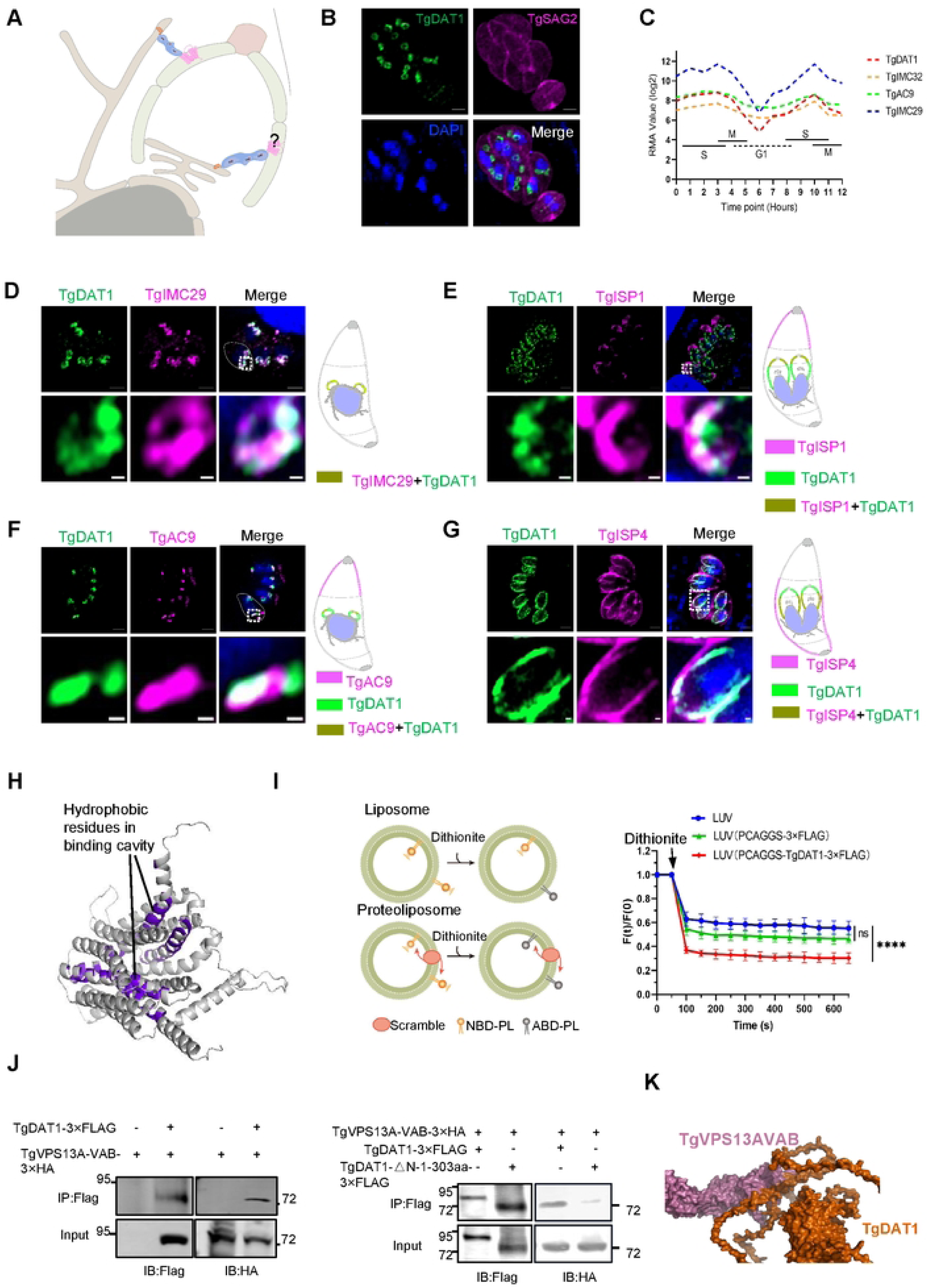
TgDAT1 is a physiologically-essential IMC-localized lipid scramblase. **(A)** A hypothetical model for Vps13-mediated lipid trafficking to IMCs involving a scramblase. **(B)** IFA showing the localization of TgDAT1. This strain was characterized by an endogenous insertion of a 3×HA tag at the C-terminus of TgDAT1. **(C)** Robust Multi-array Average (RMA) analysis of the expression of the indicated genes is presented across the cell cycle. **(D-G)** IFA analysis of the subcellular localization of TgDAT1 was performed by co-staining with TgIMC29, TgAC9, TgISP1 and TgISP4. Green: rabbit anti-HA; Magenta: mouse anti-MYC and mouse anti-TgSAG2 polyclonal antibodies. Scale bars: 2 μm (merged panels) and 0.2 μm (zoomed panels). **(H)** Surface models of TgDAT1 predicted by Alphafold 3, and the PYMOL was used to highlight the hydrophobic residues in binding cavity (purple). **(I)** A schematic diagram of the dithionite assay. Examination of the scramblase activity of TgDAT1 was conducted through the dithionite assay, with LUV and PCAGGS-3×FLAG serving as a control. Data are presented as the fluorescence intensity (FI) at indicated time points, normalized to the initial FI. The data were statistically analyzed using the Wilcoxon Signed Rank Test; ****P<0.0001. LUV represents large unilamellar vesicles without protein. **(J)** Co-immunoprecipitation assays (Co-IPs) of 3×HA-tagged TgVPS13A-VAB with 3×FLAG-tagged full-length or N terminal-deleted (1-303aa) version of TgDAT1. Cells were transiently transfected with plasmids expressing 3×FLAG-tagged TgVPS13A-VAB and 3×HA-tagged TgDAT1 or its mutant. Surface view of HADDOCK modelled complex between TgVPS13A-VAB and TgDAT1 showing the top ranked model of the best scoring cluster with TgVPS13A-VAB in magenta and TgDAT1 in vermillion color is shown between the western blotting images.

To test the interaction of TgVPS13A with TgDAT1, we co-expressed the predicted VAB domains of TgVPS13A and TgDAT1 in HEK293T cells and performed co-immunoprecipitation and insilico docking of repective predicted structures with HADDOCK. The 3xHA-tagged C-terminal VAB domain of TgVPS13A could be co-immunoprecipitated with TgDAT1-3xFLAG (Fig 5J). Our extended assays showed that the N-terminal of TgDAT1 is required for its binding with the VAB domain of TgVPS13A (Fig 5J). The TgVPS13A-VAB and TgDAT1 complex model docked using HADDOCK has a high-ranking HADDOCK score, suggesting TgDAT1 serving as a receptor for TgVPS13A in the IMC (Fig 5K).

Our structure modeling of the entire complex considering experimental results revealed TgVPS13A forming a bridge sandwiched with TgVAP and TgDAT1 proteins (Fig S5A). To further confirm the interaction of TgVPS13A, TgVAP and TgDAT1 proteins, we inserted a 4×Myc and an EGFP tag at the C-terminal of TgDAT1 or N-terminal of TgVAP respectively by CRISPR/Cas9 method. Immunostaining disclosed co-localization of TgVPS13A, TgVAP and TgDAT1 in the daughter IMC, indicating the existence of three proteins in a complex (Fig S5B-D).

### TgDAT1 plays a vital role in biogenesis of budding IMC

To assess the physiological roles of TgDAT1 in *T. gondii*, we endogenously tagged a 12×HA-AID* at the N-terminal of TgDAT1 (Fig S6A) and confirmed the correct integration of the tag through diagnostic PCRs and sequencing analysis (Fig 6A). The addition of 3-indoleacetic acid (IAA) resulted in efficient degradation of the protein, as indicated by IFA (Fig S6B) and western blotting (Fig 6B). We found that depletion of TgDAT1 inhibited the lytic cycle of parasites, as evidenced by the absence of plaque formation in IAA-treated parasites within HFF monolayers (Fig 6C).

**Fig 6.**
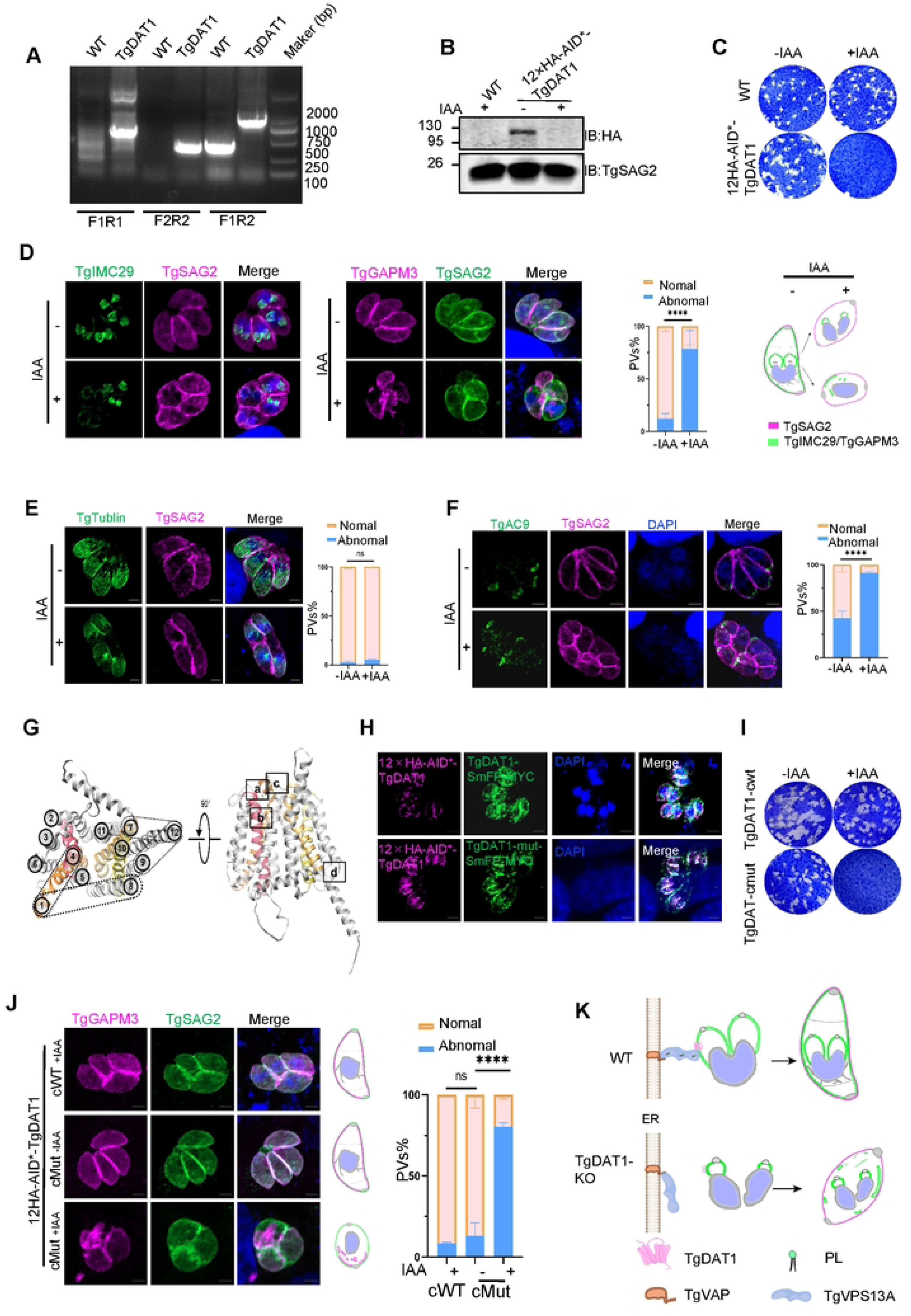
TgDAT1 plays a vital role in biogenesis of budding IMC. **(A)** PCR identification of the 12×HA-AID* tag inserted at the N-terminal of TgDAT1. **(B)** Expression of TgDAT1 was examined using western blot analysis. The parasites were cultured in the presence of IAA for 24 h in BJ-5ta cells and then subjected to the analysis. WT refers to the wild type. TgDAT1 expression was detected using anti-HA antibodies. **(C)** Plaque assays were performed by infecting BJ-5ta cells with the parasites for 9 days in the presence or absence of IAA. **(D)** The biogenesis of the daughter IMC was investigated by examining the localization of TgIMC29 and TgGAPM3 in TgDAT1-deficient parasites. An EGFP tag or a mCherry tag was inserted at the C-terminal of TgIMC29 or TgGAPM3 in the 12×HA-AID*-TgDAT1 cell line respctively. Parasites were treated with IAA for 48 h. The abnormal and normal PVs were counted from TgGAPM3. The data were analyzed for statistical significance by the unpaired t-test; ****P<0.0001. Schematic diagram of the assembly of early buds of daughter IMCs in the TgDAT1-KO parssites. **(E)** IFA analysis showing the assembly of TgTubulin in TgDAT1-deficient parasites. The abnormal and normal PVs were counted. The data were analyzed for statistical significance by the unpaired t-test; ns. **(F)** The biogenesis of the early IMC was investigated by staining with localization of TgAC9 in TgDAT1-deletion strain. Parasites were transfected with plasmids expressing TgAC9-3×MYC under the control of the the GRA1 promoter. The abnormal and normal PVs were counted. The data were analyzed for statistical significance by the unpaired t-test; ****P<0.0001. **(G)** A model of TgDAT1 and the potential sites of salt bridges predicted by AlphaFold and Chimera. **(H)** IFA analysis showing the localization of TgDAT1 mutants at the IMC. Parasites were transfected with plasmids expressing mutated versions of TgDAT1-SmFP-MYC under the control of the native promoter, and those stably expressing these mutants were selected using pyrimethamine. **(I)** Plaque assays were performed by infecting BJ-5ta cells with TgDAT1 mutants and wild-type parasites for 8 days, both in the presence and absence of IAA. **(J)** IFA showing the localization of TgGAPM3 in salt-bridge mutants of TgDAT1. An endogenous mCherry tag was inserted into the C-terminus of TgGAPM3 in parasites. The abnormal and normal PVs were counted. The data were analyzed for statistical significance by the unpaired t-test; ****P<0.0001. **(K)** A model indicating abnormal IMC biogenesis in TgDAT1-deficient parasites. Magenta: rabbit anti-TgSAG2 polyclonal antibodies, mCherry, rabbit anti-HA; Green: EGFP signal, rabbit anti-TgSAG2, rabbit anti-TgTubulin polyclonal antibodies, and mouse anti-MYC; Blue: DAPI. Scale bars: 2 μm.

Next, we investigated whether the IMC assembly was affected by the depletion of TgDAT1 by staining the nascently-recruited IMC proteins. Again, the absence of TgDAT1 resulted in stunted development of the IMC in daughter cells visualized by TgIMC29 or TgGAPM3 (Fig 6D), which resembled the TgVPS13A or TgVAP-deficient parasites. Our findings also revealed that the assembly of the cytoskeleton was not affected significantly (Fig 6E and Fig S6C). Since the apical cap may form *via* a different mechanism, we stained AC9 (a cap alveolin) and found its localization was disrupted (Fig 6F). To further determine whether the growth-stunted daughter buds in the TgDAT1-depleted parasite comprise the cap region, we colocalized TgIMC29 with AC9. While the AC9 localization pattern was primarily disrupted, its staining was still visible in the growth-stunted buds. The signal for AC9 marked only the top region of the stunted buds but did not completely merge with TgIMC29 (Fig S6D).

The conformational dynamics of transporters, including the inward-facing (IF) and outward-facing (OF) transitions, are crucial for their substrate transport functions (46). In MFS family proteins, these states may be stabilized by the formation and breaking of inter- and intra-domain salt bridges (43). The predicted structure of TgDAT1 (Fig 6G) revealed 8 residues forming salt bridges. We mutated 7 residues (316D/N, 326E/Q, 397R/Q, 576K/C, 579K/C, 617R/Q, 722E/Q) and expressed the eventual TgDAT1 mutant in the 12xHA-AID*-TgDAT1 strain. Immunostaining showed that the TgDAT1 mutant could still be positioned in the IMC (Fig 6H), but it failed to rescue the parasite growth (Fig 6I) and IMC assembly in TgDAT1-depleted strain (Fig 6J). Collectively, we show that TgDAT1 is an early-recruited IMC protein that is required for the daughter IMC assembly (Fig 6K).

### TgDAT1 is required for the phospholipid targeting to and building of the alveoli

In final assays, we examined whether membrane lipid bilayer is indeed affected upon protein depletion, leading to disrupted IMC assembly. We expressed GFP-Lact-C2 in the TgDAT1 and TgVPS13A mutants (Fig 7). GFP-Lact-C2 is a biosensor of phosphatidylserine (PtdSer) (47) and has been adopted in *T. gondii* tachyzoites to monitor the subcellular localization of PtdSer and its naturally abundant structural homolog phosphatidylthreonine (PtdThr) (48) (49). We tested localization dynamics of GFP-Lact-C2 and found that it was primarily present in the IMC of TgGAPM3-stained mature parasites, but not in the early progeny (Fig 7A). The biosensor also partially accumulated at the ER arriving sites (Fig 7B). Upon TgDAT1 depletion, GFP-Lact-C2 signal collapsed similar to that of TgGAPM3 (Fig 7C-D). We also examined its localization in the TgVPS13A-deficient parasite. Again, it was evidently mis-located and failed to reflect normal IMC (Fig 7E). The data infer a role of TgDAT1 and TgVPS13A in normal lipid expression in the alveoli membrane (Fig 7F).

**Fig 7.**
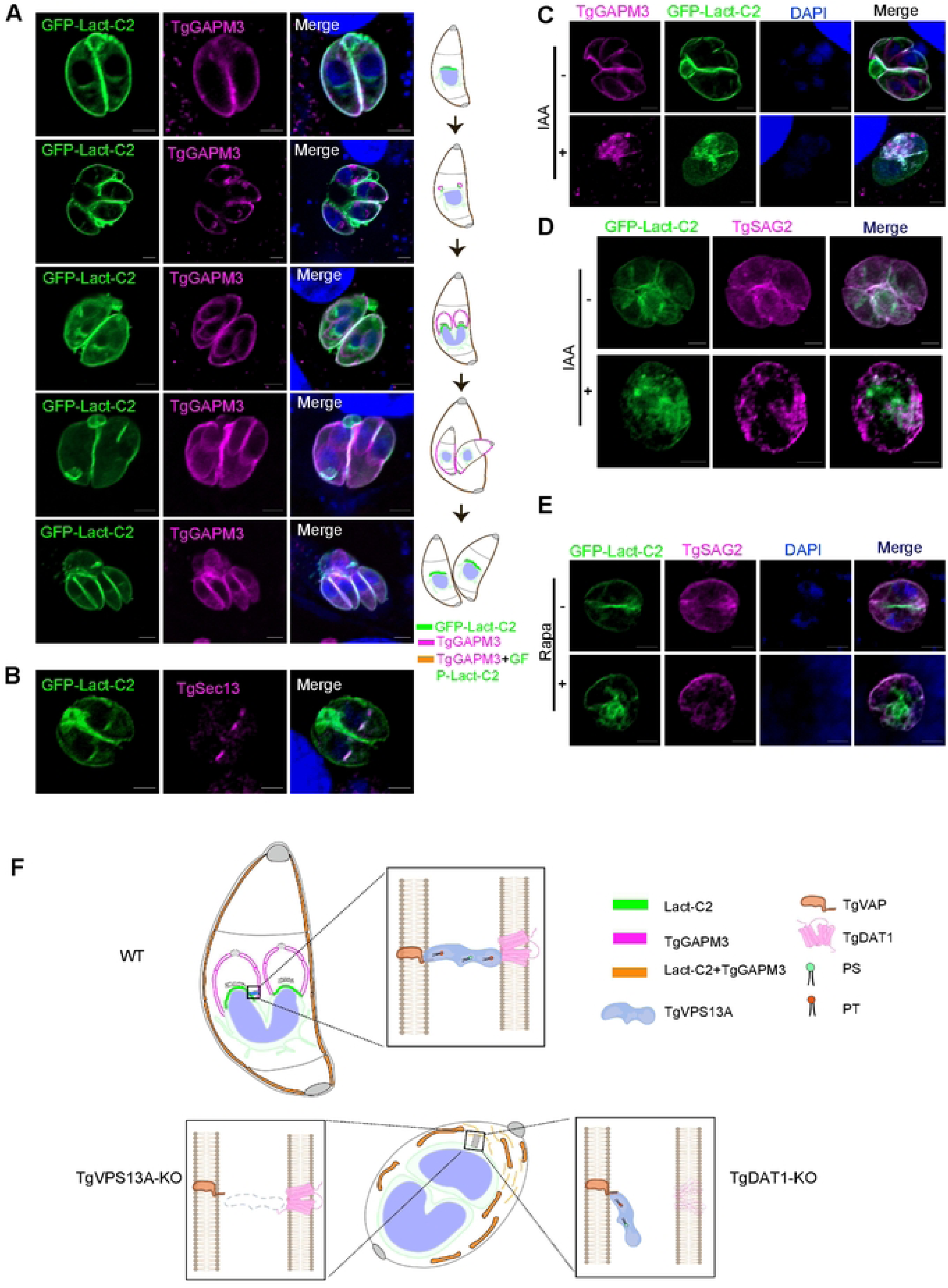
TgDAT1 is required for the phospholipid targeting to and building of the alveoli. **(A)** IFA showing colocalization of Lact-C2 and TgGAPM3 in IMCs during the complete Toxoplasma cycle. The TgDAT1-deficient strain with the mCheery inserted into the C-terminus of TgGAPM3 was transfected with a plasmid expressing GFP-Lact-C2, which is expressed under the GRA1 promoter, and those stably expressing strains were selected using pyrimethamine. **(B)** IFA analysis showing colocalization of Lact-C2 and the ER arrival sites (TgSec13) in *T. gondii*. Endogenous insertion of a 3×V5 tag into the C-terminus of TgSec13 in parasites stably expressing GFP-Lact-C2. **(C** and **D)** IFA analysis showing colocalization of LACT-C2 and TgGAPM3 in TgDAT1-deficient parasites. **(E)** IFA analysis revealing the localization of GFP-Lact-C2 in TgVPS13A-deficient parasites. **(F)** A schematic representation of PS/PT trafficking in TgDAT1- and TgVPS13A-deficient parasites. Magenta: rabbit anti-TgSAG2 polyclonal antibodies, mCheery signal and mouse anti-V5. Green: EGFP signal; Blue: DAPI. Scale bars: 2 μm.

## Discussion

Inner membrane complex is a hallmark of alveolate organisms. Using the well-established model parasitic alveolate, *Toxoplasma gondii*, we demonstrate a bridge-like complex, TgVPS13A-TgVAP-TgDAT1, at the parasite IMC-ER interface, which facilitates the biogenesis of IMC. The protein complex essential for the formation of daughter cells and thereby for asexual reproduction and lytic cycle of *T. gondii*. The proximity biotinylation strategy has identified many novel proteins associated with the IMC, including alveolins (50–53), IMC Sub-compartment Proteins (ISPs) (54), ISCs and TSCs(2). Gliding-associated proteins (GAPs) are also crucial for the IMC biogenesis, but the mechanism and function are not well understood. Until now, the mechanisms of the IMC biogenesis, especially the formation of the alveoli membrane remain unclear. Our finding of a novel protein machinery driving inter-organelle lipid transport provides a new direction to membrane and organelle biogenesis in *T. gondii* and other alveolates, including clinically-relevant protozoan parasites.

It has been shown that Brefeldin A treatment disrupts the IMC formation (20, 21). Our previous study demonstrated that proteins involved in vesicular transport, including TgVPS45 (an SM protein), TgStx6 (a Qc SNARE) (55), and Rab11A (a GTPase) (55), are necessary for the biogenesis of IMCs. Depletion of any of these proteins leads to the collapse of nascent IMCs and significant morphological changes in the daughter parasites. TgVPS45 and TgStx6 are localized at the Golgi and ELC interface. These data suggest that the vesicle fusion pathway through Golgi is required for the IMC biogenesis. This study found that TgVPS13A binds to an adaptor at the ER (TgVAP), localizes at potential membrane contact sites (MCSs) between the nascent IMC buds and the ER of the mother or daughter parasites, and enables the IMC biogenesis and cell division. Given the role of this protein family in lipid transfer between membranes in other organisms, TgVAP-TgVPS13A-TgDAT1 complex likely mediates the ER-IMC lipid transport in *T. gondii*.

Our results indicate that TgVPS13A interacts with TgVAP through its N-terminal fragment and engages with TgDAT1 *via* the C-terminal VAB domain (Fig S5). All three are essential for the parasite survival and a conditional depletion of either protein phenocopies asynchronous development of the IMC in budding parasites. A prominent dispersed staining of TgVPS13A in *T. gondii* tachyzoites also implies colocalization with other membranes, which is consistent with other VPS13 family proteins serving as tethers for organelles and promoting formation of MCSs (8). We therefore surmise a broader role of TgVPS13A in *T. gondii*. Equally, TgVPS13A-defecient parasites exhibited defective tubulin assembly and cell division. Notably, these phenotypes were not observed upon loss of TgVAP or TgDAT1. Various factors, such as PIP gradients or lipid production, promote lipid exchange at the MCSs (3). These factors may also regulate the function of TgVPS13A at the ER-MCS interface in *T. gondii*.

Different phospholipids can be salvaged and/or synthesized *de novo* (*34, 56*). PtdCho and PtdEtn, the most abundant phospholipids in *T. gondii* are made in the ER (25, 31, 57). PtdEtn in *T. gondii* can also be produced from PtdSer by TgPSD1mt in the mitochondria (26) and TgPSD1pv in the parasitophorous vacuole (58). PtdSer and PtdThr are generated in the ER (27, 32), and PtdIns in the Golgi apparatus (30), while phosphatidylglycerol (PtdGro) is thought to be produced in the ER and mitochondrion (28, 29). These major lipids must transfer from the site of *de novo* synthesis or salvage to various organelles to support the organelle biogenesis. Little is known about the subcellular distribution of distinct lipids in different organelles. Recent work of Konishi et al. deployed quick-freeze-fracture replica labeling and electron microscopy to determine the expression of PtdSer and PtdEtn in the membrane leaflets of tachyzoites (59). Notably, both lipids are primarily present in the luminal leaflet of the IMC. GFP-C2-Lact-detected phospholipids, *i.e.*, PtdSer and PtdThr, are also enriched in the IMC besides the ER (48, 49). TgVPS13A, therefore, is likely involved in transport of ER-synthesized lipids to the IMC.

It has also been demonstrated that the IMCs of mother parasites are recycled for use in daughter cells (21). We observed that the phenotype resulting from the depletion of TgDAT1 only became apparent after 48 h of treatment with IAA. TgIMC29 is a protein that primarily appears in daughter cells. We found that the development of some daughter buds was inhibited, as evidenced by the staining of TgIMC29, in parasites lacking either TgVPS13A, TgVAP, or TgDAT1. However, other daughter cells appeared to grow normally within the same vacuole. We speculate that the observed phenotype in IMC may partly be caused by the supplementation of materials derived from the mother. Previous studies in other models have shown that MCSs exist between almost all opposing membranes to facilitate lipid exchange among organelles (60–62). It is reasonable to assume that *T. gondii* may also have additional MCSs between organelles to mediate lipid transport. In this study, four putative VPS13 genes were identified in the genome of *T. gondii*. However, it remains to be determined whether other VPS13 genes and MCSs might also play a role in inter-organelle movement of lipids. Future work should investigate the roles of the remaining candidates (TGGT1_306020, TGGT1_291180, TGGT1_232080) *in T. gondii*.

## Materials and Methods

### Cell and Parasite strains culture

The hTERT-immortalized HFF cell line BJ-5ta (ATCC CRL-4001) was grown in Dulbecco’s Modified Eagle’s Medium (DMEM, Sigma-Aldrich, D6429) supplemented with 10% fetal bovine serum (CLARK, FB25015), 20% (v/v) Medium 199 basic (gibco, C11150500BT), and 1% penicillin/streptavidin. *T. gondii* RHΔhxgprtΔku80TIR1-FLAG were cultivated in BJ-5ta cells.

### Construction of plasmids

All primers used in this study are listed in Table S3. The guide RNA sequences specific for TgVPS13A (TGGT1_291180) are as follows: AATCGCCAGAGTCTCAGCGA, ATCACGCTCACGACCGCTGG, and CCAACTGCCTAACTCGACCA. For TgVAP (TGGT1_318160), the sequence is TGATTGAAACGCGAAAATGG. The sequence for TgDAT1 (TGGT1_258700) is GGCGAAGGTCTCCATTTCAT. For TgIMC29 (TGGT1_243200), the sequence is CCTTTAATTGAGGCCGTGTC. For TgGAPM3 (TGGT1_271970), the sequence is CTAAGGGACAAGGTTGACAC. For TgSec61β (TGGT1_211040), the sequence is AGAGGAGCAGAGCTCGAAGT. For TgSec13 (TGGT1_201700), the sequence is TCTCTGGAAGAGCAGAACGG. These sequences were designed using EuPaGDT (http://grna.ctegd.uga.edu/) and inserted into the PmeI site of the pCD-Cas9 vector (63) to construct CRISPR/Cas9 plasmids. The pCD-Cas9 vector contains the Cas9 gene and the TgU6 promoter, which drives the expression of the guide RNA.

The fragments for TgGAPM3-BirA*-3HA, TgSAG1-EGFP-HDEL, TgISP4-3×MYC, TgISP1-3×MYC, TgAC9-3×MYC, TgIMC29-3×MYC, and GFP-Lact-C2 were introduced into the pBluescript-DHFR vector under the control of either the the native, TgGRA1 or Tgβ-tubulin promoter. To express TgVPS13A-VAB-ATG2-C, TgVPS13A-N1-1392 and TgDAT1 in *T. gondii*, we amplified the promoter sequences and fused them with the coding sequences, a SmFP-MYC tag, and then cloned them into the pBluescript vector containing a DHFR cassette. For the TgDAT1 mutant, acidic or basic amino acids D316, E326, R397, K576, K579, R617, and E722 were replaced with neutral amino acids using overlapping PCR, and this was subsequently cloned into the aforementioned vector. To express TgVAP, TgVPS13A-N929-1392, TgVPS13A-VAB, TgDAT1 and TgDAT1-△ N-1-303 in HEK293T cells. The open reading frames of these genes were amplified, and then introduced into the PCAGGS-3×HA/PCAGGS-3×FLAG vector.

### Generation of transgenic *T. gondii* strains

To construct AID-inducible conditional knockout strains, the RHΔhxgprtΔku80TIR1-3×FLAG strain was co-transfected with a linearized DNA fragment and the pCD-Cas9 plasmid to insert the 12×HA-AID* tag at the endogenous genomic loci of TgVAP and TgDAT1 at the N-terminus. To create LoxP sites in the genome locus of TgVPS13A, we amplified the promoter region and a fragment of the N-terminal of TgVPS13A from the genomic DNA of the parasites. This amplification was flanked by two LoxP sites using PCR. The resulting construct was then used to transfect DiCre parasites with CRISPR/Cas9 plasmids to develop the conditional knockout strain. To endogenously insert tags into the C-terminus of TgVPS13A, TgIMC29, TgGAPM3, TgSec61β, and TgSec13, parasites were co-transfected with the sequences containing the SmFP-HA, EGFP and mCherry tags and pCD-Cas9 plasmid.

### Parasite lines, transfections and selection

To obtain stable above transgenic parasites, 1×10^7^ of fresh RHΔhxgprtΔku80TIR1-3×FLAG or Dicre strains were transfected with 30 μg plasmid and 6μg linearized DNA fragment by electroporation using standard procedures at 1.5 kV, 50Ω, 25 μF and 2 mm with a BTX electroporator (BIO-RAD). These parasites were allowed to grow in Vero cells for 10-16 h and then sorted with flow cytometry (SONY-SH800S). The sorted cells were inoculated into 96-well plates with fibroblast cell monolayers. After 7-8 days, monoclonal was screened by PCR to confirm correct integration. To select the strain by pyrimethamine, 1×10^7^ parasites were transfected by electroporation with 20 μg of the related plasmids. After 16 h, the transfected parasites were selected using pyrimethamine over three passages.

### Reagents and antibodies

500 mM IAA (Aladdin, I101074) was prepared with absolute ethanol. The final concentration of IAA is 500 μM. Rapamycin (MCE, HY-10219) was prepared at 10 mM in DMSO. The final concentration of Rapamycin is 50 μM. Mouse and rabbit anti-TgSAG2 (1:1000), used in immunofluorescence analysis (IFA) and western blotting (WB), were prepared previously in our lab. Tag antibodies were used at the following dilutions: mouse anti-HA mAb (Sigma-Aldrich, H9658): 1:2000 (WB), 1:1000 (IFA); mouse anti-FLAG mAb (Sigma-Aldrich, F3165): 1:2000 (WB), 1:1000 (IFA); rabbit anti-HA mAb (Cell Signaling Technology, 3724S): 1:2000 (WB), 1:1000 (IFA); mouse anti-MYC (Cell Signaling Technology, 2276S): 1:2000 (WB), 1:1000 (IFA); rabbit anti-V5 mAb(Sigma): 1:1000 (IFA). Alexa Fluor 488 secondary antibody (Invitrogen, A11001): 1:1000 (IFA); Alexa Fluor 594 secondary antibody (Invitrogen, A11037): 1:1000 (IFA); DyLight 800-labeled anti-mouse IgG (SeraCare Life Sciences, 5230-0415): 1:10000 (WB).

### Western blotting and Co-immunoprecipitation

For Western blot samples, parasites or cells was collected and incubated with RIPA buffer for 30 min on ice. Afterwards, samples were centrifuged for 30 min at 13,000 rpm at 4℃ and the supernatant was mixed with SDS loading dye (Solarbio). The samples were loaded onto a 10% SDS-PAGE gel followed by transfer onto a PVDF membrane. The PVDF membrane containing the transferred proteins was blocked with 5% skimmed milk in 1×Tris-buffered Saline (TBS) buffer containing 0.5% Tween-20 for 1 h at room temperature. The PVDF membrane was then incubated with 1:2000 dilution of rabbit anti-HA monoclonal antibody (Cell Signaling Technology, 3724S) and 1:1000 dilution of rabbit anti-TgSAG2 antibody in blocking buffer for overnight at 4°C followed by 5× washes of 1×TBST. The blots were then incubated with 1:10000 dilution of IRDyeLight 800CW goat anti-Rabbit/Mouse IgG (926-32210/926-32211) in blocking buffer for 1 h at room temperature followed by 5x washes of 1x TBST. The probed PVDF membranes were visualized using Odyssey CLX (Li-COR).

For Co-immunoprecipitation, HEK293T cells at 70% confluency in 6cm dishes were transfected with plasmids, collected after 48 h, and lysed in RAPI buffer for 30 min on ice. The supernatant (100 μL) was used for input analysis. The remaining supernatant (300 μL) was incubated with Anti-FLAG M2 Magnetic Beads (Sigma-Aldrich, M8823) or anti-HA Affinity Gel (Sigma) for overnight at 4°C with gentle rotation. The beads were washed five times with PBS and RAPI, and bound proteins were eluted with SDS-PAGE sample buffer.

### Immunofluorescence assay (IFA)

BJ-5ta monolayers grown on coverslips (CITOGLAS) or φ20 mm tissue culture dishes (Biosharp, BS-20-GJM) were infected with tachyzoites and grown at 37°C. Cells were subsequently fixed with 4% paraformaldehyde (PFA) (Biosharp, BL539A) for 30 min, and permeabilized (0.3% Triton X-100/PBS), blocked (5% BSA/0.5% Tween-20/PBS), and probed with primary antibodies 2 h prior to washing (0.5%Tween-20/PBS) and incubated with secondary antibodies (Alexa 488- or Alexa 594-conjugated goat anti-mouse/rabbit). Nuclei were stained with DAPI, and coverslips were mounted on slides (Sail brand, 7101). Image analysis was done using an LSM 980 confocal scanning microscope (Zeiss).

### Mass spectrometry of biotinylated proteins

Grow TgGAPM3-BirA*-3×HA tagged parasites in 10-cm dishes containing BJ-5ta monolayer with and without 50 µL of 160 mM D-biotin for 30h. Parasites was collected, incubated with RIPA buffer (Sigma-Aldrich, R0278) for 30 min on ice, centrifuged at 12, 000 rpm, 4 °C for 40 min. 40 μL supernatant was used to detect the expression of TgGAPM3 and TgSAG2 by WB. The other supernatant was transferred to 2 mL tubes containing 50 µL streptavidin magnetic beads (Pierce) that have been washed at 4 °C overnight. Wash beads three times with PBS, and then wash twice with RIPA buffer. The sample was boiled with loading buffer and loaded on 10% SDS-PAGE and sent to Beijing Liuhe Bada Gene Technology Co., LTD for testing.

Bands of interest were cut out and washed with ddH_2_O_2_ for 10 min. The gel pieces were then destained using an in-gel digestion and decolorization buffer (50% acetonitrile containing 25 mM NH_4_HCO_3_) for 20 min. Acetonitrile was added to dehydrate the gel pieces, which were vacuumed until they appeared completely white. Next, 10 mM dithiothreitol was added, and the gel was incubated at 56 °C for 1 h. After this, the excess dithiothreitol was removed, and the gel was incubated at room temperature in the dark with 55 mM iodoacetamide for 45 min. The excess iodoacetamide was then removed, and the gel was washed with 25 mM NH_4_HCO_3_ for 10 min. Following this wash, the gel was again treated with the in-gel digestion and decolorization buffer for 10 minutes. Acetonitrile was added for dehydration, and the gel was vacuumed until it was completely white. A trypsin solution was diluted 15 times with 25 mM NH_4_HCO_3_ and added to the dehydrated gel pieces for digestion at 37 °C overnight. The next day, 0.1% formic acid was used to stop the digestion. Finally, 10 μL of the solution containing the peptides was prepared for mass spectrometer detection.

The peptides were are separated by the UltiMate3000 RSLCnano ultra-high performance liquid system. Mobile Phase A is an aqueous solution containing 0.1% formic acid, while Mobile Phase B is an aqueous solution comprising 0.1% formic acid and 98% acetonitrile. The flow rate is maintained at 400 nL/min. The peptides are then analyzed using the Thermo Scientific™ Q Exactive™ mass spectrometer. The ion source voltage is set to 1.8 kV, and both peptide precursors and fragments are detected and analyzed using high-resolution Orbitrap technology. The mass spectrometry (MS) scanning range is set from 350 to 2000, with a scanning resolution of 70,000. Data-dependent acquisition (DDA) procedures are employed for data collection. The top 20 peptides with the highest signal intensity are selected for MS/MS analysis, using a normalized collision energy (NCE) setting of 28. Fragments are detected through heated capillary dissociation (HCD) at a resolution of 17,500.

The resulting MS/MS data were processed using MASCOT2.3.0. Tandem mass spectra were searched against ToxoDB. Trypsin was specified as cleavage enzyme allowing up to 2 missing cleavages. Mass error was set to 15 ppm for precursor ions and 20 mmu for fragment ions. Carbamidomethyl on Cys were specified as fixed modification; Gln->pyro-Glu (N-term Q), oxidation on Met and biotinylation on Lysine were specified as variable modification. Peptide ion score was set > 23. The volcano plots were created using R Studio(64) and R(65).

### Protein expression and purification

TgDAT1 was expressed in HEK293T cells for 48h. The cells were pelleted and either used immediately or stored at −20°C. For purification, the cell pellets were lysed in 10 mM HEPES PH7.4, 100 mM NaCl, 19.6mM DDM, 1x protease inhibitor cocktail using 400 µL per 10 cm dish of cells, and mixed overnight at 4 °C. Cell lysates were centrifuged at 12, 000 rpm for 40 min, and the supernatant was incubated with preequilibrated anti-FLAG M2 affinity resin (Sigma-Aldrich) for 10 h. Then, the agarose beads were washed three times in 0.5 mL of buffer W (50 mM HEPES pH 7.4, 100 mM NaCl, 1.96 mM DDM). The protein was eluted twice or three times with 10 µL per transfected dish with Buffer W supplemented with 0.2 mg/mL FLAG peptide for 8 h at 4°C with gentle mixing. Finally, the obtained target protein was quantified by BCA (Thermo).

### Dithionite assay

For preparation of liposomes and proteo-liposomes using a glass syringe, add 1, 435 µL POPC (25 mg/mL, in chloroform) and 160 µL POPG (25 mg/mL, in chloroform) to a round bottom flask to obtain 52.5 µM lipids in a molar ratio of POPC:POPG = 9:1. Then dry the lipid overnight using a rotary evaporator. Next, the resulting lipid membrane was hydrated in buffer A (10 mL of 50 mM HEPES pH 7.4, 100 mM NaCl), gently mixed for 1-2 h, and ultrasonicated for 10 min at room temperature with a frequency of 40 kHz. The liposomes were then extruded 11 times through a 400 nm pore size membrane and 5 times through a 200 nm pore size membrane. For scramblase activity assay, pipette 800 µL of LUVs into 2 mL microfuge tube, add 5.3 µL of buffer A and 34.7 µL of 10% (w/v) DDM dissolved in Buffer A, and incubate for 3 h at room temperature with end-over-end mixing. After 3 h of vesicle destabilization add the dissolved NBD-labeled phospholipid, the liposomes were incubated with purified TgDAT1 protein and 0.4 mol NBD-C6-PC (Avanti #810130P) for 1 h, and then 80 mg/mL Bio-Beads were added and incubated at room temperature for 1 h. The samples were again added with 160 mg/mL Bio-Beads and incubated at room temperature for 2 h to remove the detergent. The sample is then transferred to a new tube containing 160 mg/mL of fresh Bio-Beads. The sample is rotated overnight at 4°C. Liposomes (50 µL) containing NBD-labeled lipids were added to 1, 950 µL of buffer A and mixed. Add 40 µL of the 1 M dithionite solution (Macklin, #S817916) to the cuvette 50 s after starting the fluorescence recording and continue to record the fluorescence for a further 600 s excitation wavelength of 470 nm and emission wavelength of 530 nm by Multiscan Spectrum (Envison).

### Plaque assay

For the plaque assays, 500 parasites were inoculated into each well of 12-well plates containing a BJ-5ta monolayer and cultured with Rapa/IAA. After 24 h, the plates were supplemented with 2 mL of fresh medium, and the cells were continuously cultured for an additional 8 days. The cells were then immobilized overnight and stained with Coomassie Brilliant Blue G250 (Beyotime, ST030) for 3 h.

### Intracellular replication assay

TgVPS13A-LoxP parasites (1×10^5^) were treated with Rapa after 72 h, and were inoculated into a 12-well plate containing a BJ-5ta monolayer. After 3 h, the parasites that did not successfully invade were washed off with warm PBS. Following this, the samples were treated with Rapa for 24 h. The parasites were then fixed, stained with rabbit anti-TgSAG2 and Alexa Fluor 488-coupled anti-rabbit IgG at room temperature for 1 h and detected using IFA. Finally, the number of parasites was counted under a fluorescent microscope (Zeiss Axio Vert.A1).

### Protein structure modeling

TgVAP and TgDAT1 structures were modelled with Alphafold3(66). TgVPS13A was modeled as three distinct parts: the N terminus, the Bridge, and the C terminus, and then stitched together using PyMOL (The PyMOL Molecular Graphics System, Version 3.1 Schrödinger, LLC.) as described elsewhere(67). The stitched structure was manually curated to highlight the backbone of the protein with the lipid transfer bridge and the membrane binding domains on N and C terminus. The protein complex docking was performed by the HADDOCK 2.4(13, 68) and visualized by the Biovia Discovery Studio(69). The potential sites for salt bridge were predicted by the Chimera.

### Oligonucleotides

All primers used in this study are listed in Table EV2.

### Statistical analyses

Data were analyzed using GraphPad Prism 9 software. The results are presented as the mean ± standard deviation (SD) from triplicate, parallel, independent experiments, unless stated otherwise. All statistical analyses were conducted using either the student’s t-test or a two-way ANOVA, unless specified differently.

## Author contributions

LZ performed most of the experiments. JF conducted the screening of transporters; HC and YZ constructed the plasmid library of the transporters for screening; SP conducted the modeling and phelogenetic analysis of the complex under the supervision of NG. LZ, QJ, NG and HJ drafted the manuscript. HJ conceived and supervised the research.

## Acknowledgments

We are grateful to the *Toxoplasma* community for sharing specified resources. The pCD-Cas9 vector was modified from the pSAG1::CAS9-U6::sgUPRT vector. pSAG1::CAS9-U6::sgUPRT, pTUB1:OsTIR1-3FLAG, SAG1:CAT, pTUB1:YFP-mAID-3HA, and DHFR-TS:HXGPRT were gifts from David Sibley (Addgene plasmids #54467, #87258, and #87259). We thank Ming Wang for the help to predict the residues responsible for salt bridge formation. This work was supported by the Natural Science Foundation of Heilongjiang to HJ (no. JQ2022C006), the National Key Research and Development Program of China (no. 2022YFD1800200) to HJ and QJ, and DBT–Wellcome Trust Senior Fellowship grant to NG (India Alliance, IA/S/19/1/504263). The funders had no role in the design, data collection, analysis, or decision to publish this work.

## Supporting information captions

**Fig S1. Identification of VPS13 family proteins in *T. gondii*.**

**(A)** Diagram of VPS13 family proteins. **(B)** Alphafold-predicted and manually curated structure of TgVPS13A. **(C)** A schematic diagram of inserting a SmFP-HA tag at the C-terminal of TgVPS13A. **(D)** PCR analysis confirming the SmFP-HA insertion at the C-terminal of TgVPS13A.

**Fig S2. Identification of VAP family protein in *T. gondii*.**

**(A)** Multialignment of TgVAP with its human orthologues by Clustal Omega. **(B)** Alphafold-predicted and manually curated structure of TgVAP. **(C)** Surface view of HADDOCK modelled complex between TgVPS13A (N term 900-1500aa) and TgVAP showing the top ranked model of the best scoring cluster with TgVPS13A in orange and TgVAP in blue color.

**Fig S3. A** proximity biotinylation assay showing TgVPS13A and TgVAP are associated with the IMC. **(A)** A model of proximity biotinylation assay with TgGAPM3-BirA*. **(B)** A schematic diagram illustrating the expression cassette that encodes TgGAPM3-BirA*-3HA. IFA showing the expression and localization of TgGAPM3-BirA* in parasites with and without biotin. Magenta: rabbit anti-HA antibodies; Green: streptavidin-Alexa 488. **(C)** Western blot analysis demonstrating the presence of biotinylated proteins in lysates from TgGAPM3-BirA* parasites. **(D)** Proximity biotinylation candidate enrichment revealing several proteins in the vicinity of TgGAPM3. **(E)** Proteins associated with the IMC were identified in the TgGAPM3-BioID analysis. **(F)** IFA showing the colocalization of TgSec61β with the ER-resident protein TgSAG1-GFP-HDEL. Magenta: mouse anti-V5; Green: EGFP signal; Blue: DAPI. Scale bars: 2 μm (merged panels) and 0.2 μm (zoomed panels).

**Fig S4. A global screening of annotated transporters in ToxoDB.**

**(A)** A global screening of annotated transporters in ToxoDB was conducted by inserting a HA tag endogenously using the CRISPR/Cas9 method. **(B)** IFA showing the localization of 64 proteins.

**Fig S5 TgVAP-TgVPS13A-TgDAT1 forms a complex for lipid targeting to the nascent IMC.**

**(A)** Full length of TgVPS13A with TgVAP and TgDAT1 fitted in the interacting pockets identified with HADDOCK. **(B)** IFA showing the colocalization of TgDAT1, TgVPS13A, and TgVAP at the early and late stages of IMC budding. In TgVPS13A-SMFP-HA parasites, a 4×MYC tag or an EGFP tag was endogenously inserted into the C-terminus of TgDAT1 or the N-terminus of TgVAP respectively. Magenta: rabbit anti-HA; Green: EGFP signal; Blue: mouse anti-MYC. Scale bars: 2 μm (merged panels) and 0.2 μm (zoomed panels). **(C)** A schematic diagram showing the lipid bridges of the TgVAP-TgVPS13A-TgDAT1 complex formed at the ER-IMC interface during the early and late stages of daughter buds. **(D)** A predicted model of TgVAP-TgVPS13A-TgDAT1 forming a lipid transport bridge between the ER and IMC. The predicted model is constructed based on data presented in this study and our understanding of lipid transport in other eukaryotic systems. Three distinct projections of the TgVAP-TgVPS13A-TgDAT1 complex are depicted to visualize the protein-protein interaction and hydrophobic tunnel in TgVPS13A to facilitate the transport of lipids (exemplified herein by PtdSer and PtdThr). Embedding of the complex in the membrane of the ER and IMC is only for the illustration purpose (not modeled or simulated for lipid bilayer-protein interactions). Note that only protein complex is rotated 360°, not the membrane bilayer for simplicity. Created in BioRender. (2025) https://BioRender.com/bte8xdo

**Fig S6 Phenotypic analysis of IMC assembly in the TgDAT1-deleted parasites.**

**(A)** Schematic diagram of inserting a 12×HA-AID* tag at the N-terminal of TgDAT1. **(B)** Immunofluorescence analysis was conducted to assess TgDAT1 expression in the parasite, both in the presence and absence of Indole-3-acetic acid (IAA). The parasites were cultured with or without IAA for 24 h and stained with antibodies against TgSAG2 and HA. **(C)** The biogenesis of the IMC was investigated by staining with IMC1 antibodies in TgDAT1-deficient parasites. The parasite lines were cultured with or without IAA for 48 h and stained with antibodies against TgSAG2 and TgIMC1. **(D)** Colocalization of TgAC9 and TgIMC29 in atrophic daughter IMCs in the TgDAT1 deletion strain. A schematic diagram of TgIMC29 colocalization with TgAC9 on atrophic daughter IMCs in TgDAT1-deficient parasites is shown at the right side. Magenta: mouse anti-TgSAG2 polyclonal antibodies; Green: rabbit anti-HA antibody; Blue: DAPI. Scale bars: 2 μm.

